# DivIVA concentrates mycobacterial cell envelope assembly for initiation and stabilization of polar growth

**DOI:** 10.1101/341073

**Authors:** Emily S. Melzer, Caralyn E. Sein, James J. Chambers, M. Sloan Siegrist

## Abstract

In many model organisms, diffuse patterning of cell wall peptidoglycan synthesis by the actin homolog MreB enables the bacteria to maintain their characteristic rod shape. In *Caulobacter crescentus* and *Escherichia coli*, MreB is also required to sculpt this morphology *de novo*. Mycobacteria are rod-shaped but expand their cell wall from discrete polar or sub-polar zones. In this genus, the tropomyosin-like protein DivIVA is required for the maintenance of cell morphology. DivIVA has also been proposed to direct peptidoglycan synthesis to the tips of the mycobacterial cell. The precise nature of this regulation is unclear, as is its role in creating rod shape from scratch. We find that DivIVA localizes nascent cell wall and covalently associated mycomembrane but is dispensable for the assembly process itself. *Mycobacterium smegmatis* rendered spherical by peptidoglycan digestion or by DivIVA depletion are able to regain rod shape at the population level in the presence of DivIVA. At the single cell level, there is a close spatiotemporal correlation between DivIVA foci, rod extrusion and concentrated cell wall synthesis. Thus, although the precise mechanistic details differ from other organisms, *M. smegmatis* also establish and propagate rod shape by cytoskeleton-controlled patterning of peptidoglycan. Our data further support the emerging notion that morphology is a hardwired trait of bacterial cells.

## Introduction

Bacteria adopt a variety of characteristic shapes, each with distinct advantages. For example, rod morphology optimizes the surface area to volume ratio and may promote nutrient uptake [Young 2006]. Rod shape is maintained by synthesis of cell wall peptidoglycan [Daniel and Errington 2003], a rigid biopolymer that encases the cell and counteracts turgor pressure. Many well-studied species, including *Escherichia coli* and *Bacillus subtilis*, elongate by adding new peptidoglycan along the lateral cell body [de Pedro et al. 1997; Daniel and Errington 2003; Wang et al. 2012; Scheffers and Pinho 2005; Tiyanont et al. 2006; Liang et al. 2017]. In these organisms, cell shape and spatial regulation of peptidoglycan synthesis are both tied to the actin homolog MreB [van den Ent et al. 2001; Gitai et al. 2004; Garner et al. 2011; van Teeffelen et al. 2011; Dominguez-Escobar et al. 2011; Errington 2015]. Inhibition of MreB causes the bacteria to transform into spherical [Doi et al. 1988; Wachi et al. 1989; Jones et al. 2001; Gitai et al. 2004] or lemon-shaped cells [Takacs et al. 2010], that for *E. coli* have been shown to have active peptidoglycan metabolism [Ranjit et al. 2017; Billings et al. 2014; Margolin 2009]. MreB is one of several bacterial proteins that are now widely accepted as components of the bacterial cytoskeleton, which have been reviewed in [Cabeen and Jacobs-Wagner 2010; Errington 2015; Carballido-Lopez and Errington 2003; Eun et al. 2015; Wagstaff and Lowe 2018]. Similar to their eukaryotic homologs, polymerization of bacterial cytoskeletal proteins is important for their roles in cellular functions, including shape maintenance [Carballido-Lopez and Errington 2003; Eun et al. 2015; Wagstaff and Lowe 2018]. These proteins also exhibit structural, although not necessarily sequence similarity, to eukaryotic cytoskeleton proteins.

Most work on bacterial morphology has concentrated on defining pathways that propagate pre-existing shape. Creating or recreating a shape, however, poses distinct spatial challenges. While studying the rod-to-sphere transition can illuminate mechanisms of shape maintenance, tracking the sphere-to-rod reversion can similarly shed light on the requirements for *de novo* morphogenesis [Billings et al. 2014; Ranjit and Young 2016; Ranjit et al. 2017; Kawai et al. 2014; Takacs et al. 2010]. One way to study rod formation is to generate spheroplasts that lack a cell wall, without which bacteria swell and become spherical (Fig. 1) [Onoda et al. 1987; Udou et al. 1982; Udou et al. 1983; Birdsell and Cota-Robles 1967]. This state is reversible: not only can cells rebuild their wall, but they are also able to recover rod morphology. In *E. coli*, sphere-to-rod reversion requires MreB to direct peptidoglycan synthesis in the appropriate locations [Billings et al. 2014]. This means that the same machinery that maintains rod morphology can also drive *de novo* rod morphogenesis. Thus *E. coli* morphology is thought to be a hardwired property of the cell that depends on MreB [Billings et al. 2014].

**Figure 1.**
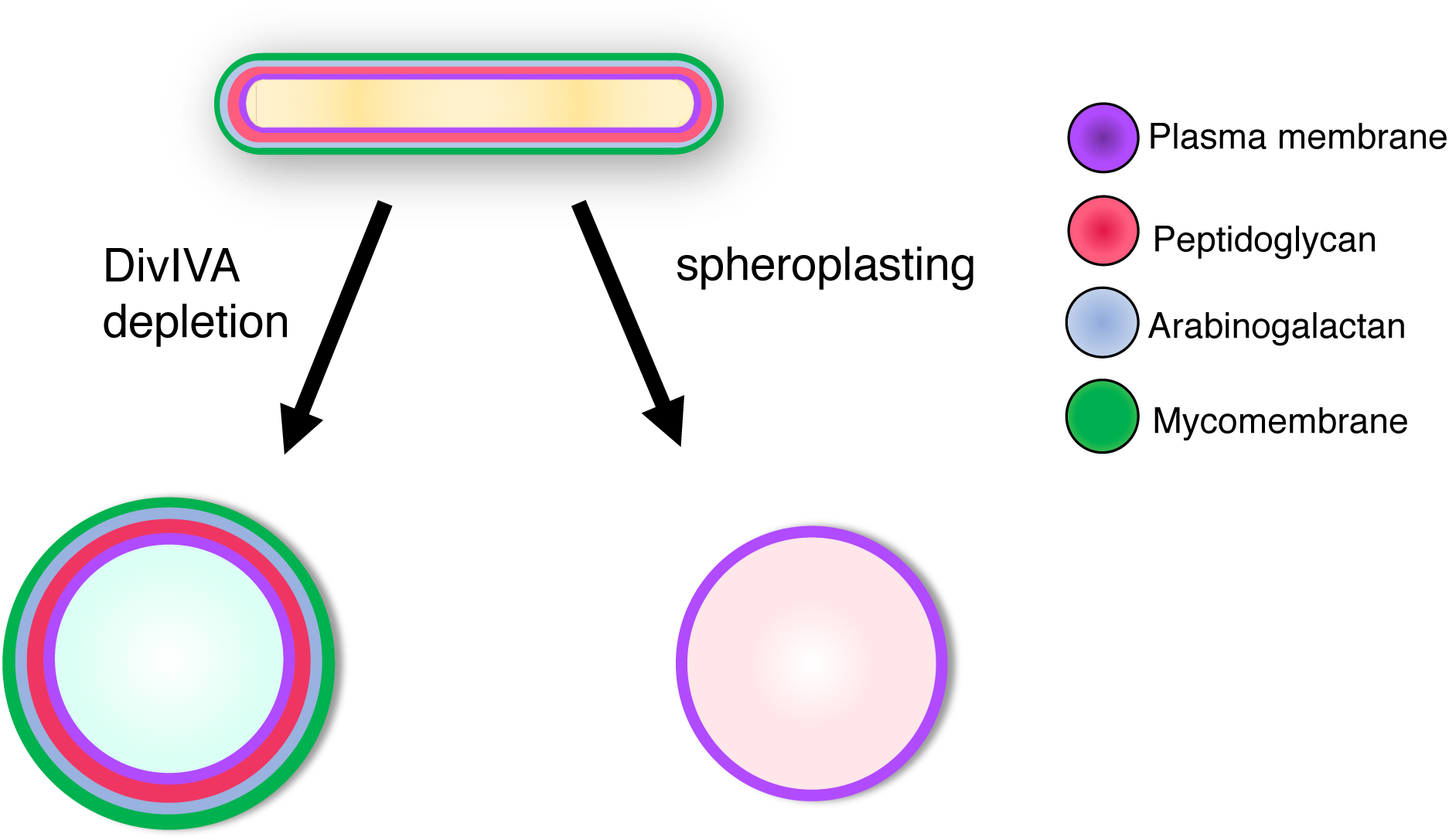
DivIVA depletion and cell wall digestion result in spherical mycobacterial cells that differ in envelope composition.

While extensive work has been done to elucidate the key role of MreB in regulation of bacterial elongation by lateral insertion of peptidoglycan, it is not the only means by which rod shaped bacteria maintain their morphology. Actinobacteria, including the pathogen *Mycobacterium tuberculosis* and the model organism *Mycobacterium smegmatis*, synthesize peptidoglycan predominantly at the cell poles [Daniel and Errington 2003; Thanky et al. 2007]. Mycobacterial elongation is asymmetric; extension of the old pole is more rapid than that of the new pole [Aldridge et al. 2012; Joyce et al. 2012; Singh et al. 2013; Rego et al. 2017; Botella et al. 2017; Meniche et al. 2014; Siegrist et al. 2015]. The mycobacterial cell envelope includes additional, covalently-attached layers that are not found in other genera, including the glycoconjugate arabinogalactan and the mycolic acids that comprise the mycomembrane [Jankute et al. 2015].

The building blocks of the mycobacterial envelope are synthesized in the cytoplasm, transported across the plasma membrane, and assembled extracellularly [Xu et al. 2017; Jankute et al. 2015]. Construction of the different layers is likely to be spatially coordinated to ensure envelope integrity and cell shape. However, mycobacteria do not encode an obvious homolog for MreB. Instead the essential protein DivIVA may fulfill this organizational role [Nguyen et al. 2007; Kang et al. 2008; Meniche et al. 2014]. DivIVA homologs are found in other bacteria but their function appears to be highly species-specific [Cha and Stewart 1997; Marston and Errington 1999; Thomaides et al. 2001; Ramirez-Arcos et al. 2005; Flardh 2003; Ramos et al. 2003]. The proteins exhibit structural homology to the eukaryotic cytoskeletal protein tropomyosin which plays a key role in yeast morphology [Drees et al. 1995; Gunning et al. 1997].

DivIVA marks the tips of mycobacterial cells, and is found in higher concentrations at the older, faster growing pole [Nguyen et al. 2007; Kang et al. 2008; Botella et al. 2017; Meniche et al. 2014]. Depletion of the protein causes dramatic morphological changes, namely the formation of spherical cells [Nguyen et al. 2007; Kang et al. 2008; Meniche et al. 2014]. These spherical cells are more likely to lyse, suggesting that the morphological changes are accompanied by decreased envelope integrity. The genomic context of *divIVA* (*wag31*) also supports a role in envelope homeostasis, as the gene is located in a cluster that also encodes many key enzymes in peptidoglycan metabolism [Kang et al. 2008]. To test whether DivIVA depletion affected peptidoglycan synthesis, the authors labeled the cells with fluorescent vancomycin, an antibiotic that preferentially marks new peptidoglycan. However, they were unable to determine whether peptidoglycan synthesis was absent or simply dispersed [Kang et al. 2008]. Phosphorylation, overexpression or tagging of DivIVA correlates with changes to the sites of peptidoglycan synthesis and cell growth [Nguyen et al. 2007; Kang et al. 2008; Meniche et al. 2014; Botella et al. 2017; Jani et al. 2010; Hamasha et al. 2010]. Although the phosphorylation status of DivIVA impacts peptidoglycan precursor synthesis, a direct, physical interaction between the synthetic enzymes and DivIVA has not been detected, making it unclear if DivIVA directly regulates their activity [Jani et al. 2010].

When DivIVA is overexpressed in mycobacteria, a subset of bacteria have ectopic, DivIVA-marked poles that branch off of the main cell body [Nguyen et al. 2007]. Overproduction of DivIVA in the distantly-related species *Streptomyces coelicolor* initiates new regions of cell wall synthesis [Hempel et al. 2008]. These observations prompted us to consider whether DivIVA, like MreB, might function in *de novo* rod morphogenesis in addition to its known role in shape maintenance. Here we show that DivIVA is required for population-wide reversion of mycobacterial spheres to rods. There is also a tight spatiotemporal association between DivIVA location and site of rod extrusion. We hypothesized that DivIVA may contribute to *de novo* morphogenesis via a mechanism similar to the one it uses to maintain existing rod shape. Using metabolic labeling paired with high resolution microscopy, we find that assembly of peptidoglycan and the mycomembrane occurs but is disorganized upon DivIVA depletion. Furthermore, repletion of the protein is required to re-localize envelope assembly. Taken together, our data suggest that DivIVA programs mycobacterial morphology by organizing envelope synthesis.

## Results and Discussion

### DivIVA contributes to de novo rod morphogenesis in mycobacteria

Depletion of DivIVA results in spherical cells [Nguyen et al. 2007; Kang et al. 2008; Meniche et al. 2014]. Sassetti and coworkers, including author MSS, found that rod morphology could not be recovered after DivIVA repletion, suggesting that DivIVA is required for shape maintenance but may not be sufficient for *de novo* rod formation. The investigators hypothesized that the spherical state is irreversible because DivIVA requires negative membrane curvature to localize and to form a new pole [Meniche et al. 2014; Lenarcic et al. 2009; Huang and Ramamurthi 2010]. However, we noted that spherical mycobacterial cells generated by cell wall digestion [Rastogi and Venkitasubramanian 1979; Yabu and Takahashi 1977; Udou et al. 1982; Udou et al. 1983], rather than DivIVA depletion, have been reported to reform rods. Spheroplasting offered a means to track *de novo* morphogenesis from spherical cells that were formed independently of DivIVA status. While elucidation of the role of MreB in bacterial morphogenesis has been greatly accelerated by the use of fast-acting small molecule inhibitors [Foss et al. 2011], to our knowledge there is currently no direct inhibitor of DivIVA [Singh et al. 2017]. Accordingly, we generated spheroplasts of the *M. smegmatis* DivIVA-eGFP inducible degradation strain used previously for depletion and repletion of the protein [Meniche et al. 2014]. This mutant expresses a single copy of *divIVA* from its native promoter. The resulting protein is C terminally tagged with eGFP and an inducible degradation (ID) tag, which allows conditional degradation of the protein upon addition of anhydrotetracycline (ATC) [Meniche et al. 2014; Wei et al. 2011]. The DivIVA-eGFP strain is viable, indicating that the tagged, essential protein is functional. We first confirmed loss of fluorescently-labeled peptidoglycan and of rod morphology following the enzymatic digestion procedure (Figs. S1 and 2B). We then allowed the cells to recover in the absence or presence of ATC. At 0, 24 and 48 hours of recovery, cells were blindly scored as one of four morphologies: sphere, early transition, late transition, or rod. The DivIVA-eGFP spheroplast population exposed to ATC was significantly slower to revert to rod morphology than the one that was not (Fig. 2A, 2B). By contrast, there was no ATC-dependent difference in population outgrowth when the spheroplasts were generated from wildtype *M. smegmatis*.

**Figure 2.**
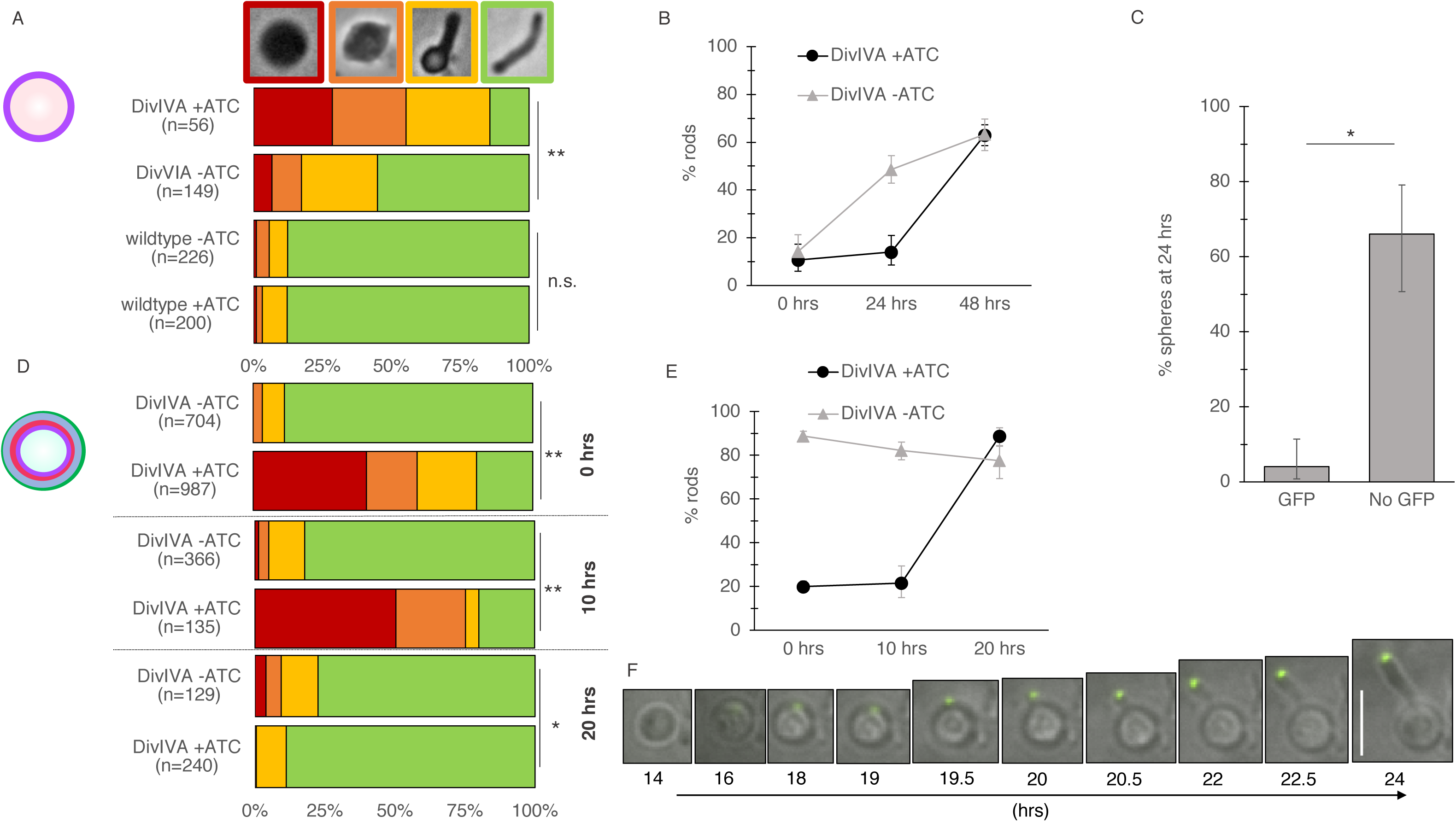
DivIVA contributes to *de novo* rod formation in *M. smegmatis*. (A) Percentage of cells in the population that exhibit a given morphology 24 hours after spheroplasting. Shape exemplars are in red, orange, yellow and green boxes above. Wildtype or DivIVA-eGFP *M. smegmatis* were converted to spheroplasts then recovered in the presence or absence of ATC for 24 hours. Cell morphology was blindly scored. Data are representative of 3 independent biological replicates. **, p < 0.01 two-tailed z test for population proportions. (B) Percentage of rod shaped cells over time post-spheroplasting, 131 <n<490. Error bars, 95% confidence intervals. (C) Proportion of eGFP-positive (n=74) and eGFP-negative cells (n=47) with spherical morphology 24 hours recovery post-spheroplasting. DivIVA-eGFP strain was incubated in ATC during outgrowth. Error bars, 95% confidence intervals. Data are representative of two independent biological replicates. (D) Percentage of cells in the population that exhibit a given morphology after DivIVA repletion. DivIVA-eGFP was incubated in the presence of ATC for 16 hours. Cell morphology was blindly scored after washout of ATC. 129<n<987. *, p < 0.05, **, p < 0.001, two-tailed z test for population proportions. (E) Percentage of rod shaped cells over time post-spheroplasting, 129<n<987. Error bars, 95% confidence intervals. (F) Time-lapse microscopy tracking of a cell recovering from 16 hours DivIVA-eGFP depletion, starting 14 hours post-ATC removal, representative of 21 cells. Scale bar: 5 µm.

We considered two, non-mutually exclusive explanations for the observation that rod formation was delayed but not blocked in the presence of ATC. First, DivIVA might contribute to, but not be required for, sphere-to-rod transition at the population level. Alternatively, or additionally, degradation of ATC or the appearance of ATC-insensitive escape mutants may have allowed a DivIVA-eGFP-positive subpopulation to outcompete the parental genotype and reform rods. To distinguish between these possibilities, we took advantage of the fact that DivIVA is tagged with eGFP and thus easily detected in single cells. We first asked whether ATC addition was associated with eGFP absence. While there is generally a close, inverse correlation between ATC and eGFP at early depletion time points, at 24 hours only 39% of ATC-treated bacteria (n=121) remained eGFP-negative (compared to 4% of untreated bacteria, n=271). If DivIVA is indeed required for the sphere-to-rod transition, we hypothesized that, regardless of the underlying cause of DivIVA repletion in the presence of ATC, preservation of spherical morphology should only occur when DivIVA-eGFP is absent. Indeed, we find that eGFP-negative bacilli are significantly more likely than their eGFP-positive counterparts to remain spherical at 24 hours (Fig. 2C). The close correlation between the absence of DivIVA-eGFP and the apparent retention of spherical morphology raised the possibility that DivIVA is required for creation of cell shape in addition to the maintenance of this phenotype.

In contrast to DivIVA in *B. subtilis* protoplasts [Ramamurthi and Losick 2009], a single focus of mycobacterial DivIVA-eGFP appeared to remain associated with the cell surface after spheroplasting, in the absence of visible negative curvature (Fig. S2). This was also true of an N-terminally tagged RFP-DivIVA fusion protein that was expressed under a strong, heterologous promoter in a merodiploid strain (Fig. S2). We wondered whether the preexisting focus of DivIVA-eGFP might facilitate rod formation and, therefore, explain the apparent discrepancy in *M. smegmatis* ability to reform rods after spheroplasting (Fig. 2B) but not after DivIVA depletion [Meniche et al. 2014]. We also considered the possibility that the difference in phenotype may be due to the different time frames of the two experiments; whereas spheroplasts required 24-48 hours to regenerate a population of rod-shaped cells [Udou et al. 1982] (Fig. 2B), Sassetti and coworkers investigated the effects of DivIVA repletion after 12 hours. To resolve this discrepancy, we revisited the question of whether DivIVA-depleted spheres could, with more time, reform rods upon ATC removal. We grew the DivIVA-eGFP strain in the presence or absence of ATC and again blindly scored individual cells for morphology. As expected, addition of ATC resulted in the appearance of bulged and spherical cells after 16 hours (designated as time 0 hour, Fig. 2D, 2E). We then washed away the ATC to block the depletion of the protein. The proportion of rod-shaped cells did not appreciably change up to 10 hours after ATC removal, a finding that was again consistent with published work. However, an additional 10 hours of growth in the absence of ATC resulted in a near-uniform population of rod-shaped cells (Fig. 2D, 2E). During both depletion and recovery, ATC-treated cells grew more slowly than their non-treated counterparts (Fig. S3).

Whether spherical mycobacteria were generated by cell wall digestion or by depletion of DivIVA, our data suggested that the presence of the protein correlated with population-wide transition to rod shape. Despite our efforts to obtain a homogenous starting population of spheres, we noted that 5-20% of cells retain rod shape following enzymatic cell wall digestion or DivIVA depletion. We considered the possibility that outgrowth of this subpopulation might account for the population-wide shift from spherical cells to rods, rather than shape alterations of individual cells. In this scenario, the spherical state is irreversible, either because such cells are not viable or because they are unable to regain rod morphology. The doubling time of the DivIVA-eGFP strain is ~4 hours, slightly slower than wildtype *M. smegmatis* [Meniche et al. 2014] (Fig. S4). The 10 or 24 hour window for shape recovery following DivIVA depletion or spheroplasting, respectively, left open the possibility that the sphere-to-rod transitions occur at the population level but not in single cells. Accordingly, we used time-lapse microscopy to track morphogenesis in single DivIVA-eGFP cells rendered spherical by ATC incubation. After ATC washout, the protein first appeared dispersed throughout the cell at ~14 hours then coalesced into a single focus. A new, DivIVA-marked pole protruded from this site (Fig. 2F, Movie S1). Organisms whose shape is maintained by MreB have been shown to achieve *de novo* rod morphogenesis by different strategies. Gradual cell thinning and elongation, discrete polar protrusion, and division prior to rod regeneration have all been observed in sphere-to-rod transition of such bacteria [Ranjit et al. 2017; Takacs et al. 2010; Billings et al. 2014; Hussain et al. 2018]. However, we observed that mycobacterial DivIVA-mediated sphere-to-rod reversion is only accomplished via protrusion of a new localized growth pole. Such protrusions occurred in 37% of the population (n=52). None of the remaining 63% of spherical cells exhibited visible eGFP foci (Fig. S5). Therefore, we hypothesized that at least some of the failure to generate rod shape was due to cell death, rather than the inability of DivIVA to initiate polar growth. We tested the viability of DivIVA-depleted *M. smegmatis* with propidium iodide, which only penetrates cells with compromised plasma membranes. Indeed we found that 36% of DivIVA-depleted spherical cells (n=94) were nonviable (Fig. S5). This finding explains approximately half of the failure to recover. The lack of recovery in the remaining, viable subset of spheres may be from slow turnover of the introduced, ATC-controlled HIV protease [Wei et al. 2011; Meniche et al. 2014] in non-dividing cells, which in turn would depress the levels of DivIVA-eGFP. These data indicate that spherical mycobacteria are able to recreate rod shape, and furthermore, that there is a close spatial and temporal correlation between DivIVA and *de novo* pole formation.

DivIVA has been hypothesized to localize based on the negative curvature of the membrane [Meniche et al. 2014] as it does in *B. subtilis* [Lenarcic et al. 2009; Huang and Ramamurthi 2010]. Unlike its *B. subtilis* homolog, however, mycobacterial DivIVA does not directly bind the membrane, suggesting that its polar localization might be achieved by a different mechanism [Plocinska et al. 2012; Plocinski et al. 2013; Plocinski et al. 2011]. We find that DivIVA is able to form a single focus in spherical cells, and that that site is also the site of rod extrusion (Fig. 2F, Movie S1). Fluorescent DivIVA foci persist upon spheroplasting (Fig. S2). Taken together, these observations suggest that negative curvature may not be required for recruitment or for stabilization of the DivIVA oligomer at the membrane.

### DivIVA is required for localized cell envelope assembly in mycobacteria

How might DivIVA contribute to morphogenesis? In mycobacteria, synthesis of cell envelope components, including peptidoglycan, arabinogalactan and the mycomembrane, occurs predominantly at the poles, as indicated by location of both the synthetic enzymes and their products [Meniche et al. 2014; Foley et al. 2016; Swarts et al. 2012; Botella et al. 2017; Thanky et al. 2007; Aldridge et al. 2012; Hayashi et al. 2018]. Several lines of evidence suggest that DivIVA is key for orchestrating envelope synthesis at the tips of the cell. Phosphorylation of the protein positively regulates its localization and correlates with more intense polar staining by fluorescent vancomycin [Jani et al. 2010]. DivIVA phosphorylation status also correlates with the capacity of isolated membrane fractions to support peptidoglycan precursor synthesis from radiolabeled substrate [Jani et al. 2010]. Overexpression of a fluorescent DivIVA fusion protein alters the interpolar distribution of both the protein itself as well as metabolic labeling of peptidoglycan [Meniche et al. 2014; Botella et al. 2017]. However, a previous study was unable to determine whether new peptidoglycan was absent or simply dispersed when the protein was depleted [Kang et al. 2008]. While the former might indicate that DivIVA regulates the activity of peptidoglycan synthetic enzymes, the latter could suggest that DivIVA controls their localization. Distinguishing between these possibilities is critical for understanding how DivIVA orchestrates cell envelope assembly and morphogenesis.

We sought to clarify the nature of DivIVA regulation by improving the spatial and temporal precision of nascent envelope detection via metabolic labeling and high-resolution imaging. In contrast to staining by antibiotic conjugates, labeling by tagged precursors of peptidoglycan or mycomembrane can reveal the presence of nascent envelope within minutes [Siegrist et al. 2015; Siegrist et al. 2013; Foley et al. 2016] and does not obviously interfere with bacterial growth [Siegrist et al. 2015]. The peptide portion of the peptidoglycan biopolymer terminates in two D-alanine residues. Derivatives of D-alanine and D-alanine-D-alanine have been used in many bacterial species, including mycobacteria, to detect cell wall metabolism [Siegrist et al. 2013; Kuru et al. 2012; Liechti et al. 2014; Lebar et al. 2014; Fura et al. 2015; Siegrist et al. 2015; Meniche et al. 2014; Botella et al. 2017; Hayashi et al. 2018]. Single residue D-amino acid probes have been hypothesized to incorporate into peptidoglycan in part or in whole via periplasmic transpeptidases [Siegrist et al. 2015]. D-alanine-D-alanine dipeptide probes are likely to be integrated into peptidoglycan at an earlier, cytoplasmic step [Liechti et al. 2014; Sarkar et al. 2016]. Together the two classes of probes can be used to track both remodeling and synthesis of the biopolymer.

We examined the effect of DivIVA depletion on peptidoglycan assembly by labeling cells with an alkyne-bearing dipeptide probe (alkDADA) [Liechti et al. 2014], which was detected by copper-catalyzed azide-alkyne cycloaddition (CuAAC) [Siegrist et al. 2015] to picolyl azide-TAMRA, or with a single residue D-alanine probe conjugated directly to the TAMRA dye (RADA) [Kuru et al. 2012; Kuru et al. 2015]. To increase our confidence in the spatial details of labeling, we imaged the bacteria using Structured Illumination Microscopy (SIM). We incubated our DivIVA-eGFP strain in the presence or absence of ATC and added alkDADA or RADA for the final 15 min (~10% generation) of culture time. In the absence of ATC, fluorescent signal derived from the probes localized primarily to the septa and DivIVA-eGFP-marked poles, with limited sidewall labeling (Fig. 3A, 3C). After 4 hours of ATC incubation, most cells were eGFP-negative and misshapen. The morphological changes were accompanied by dramatic alterations in labeling: TAMRA fluorescence was no longer polar, and either partially or completely surrounded the cell (Fig. 3B, 3D). After 8 hours of depletion, the cells developed large bulges that were evenly labeled, and after 12 hours, they were primarily spherical, bounded by homogenous labeling (Fig. 3B, 3D). Some of the eGFP-negative spheres had labeled septa (Fig. 3B), supporting the idea that the divisome is regulated independently of DivIVA [Kang et al. 2008; Santi et al. 2013]. Continued incorporation of alkDADA in the absence of DivIVA suggested that in addition to periplasmic remodeling of peptidoglycan, cytoplasmic synthesis also persists. To further distinguish between the two processes, we repeated the labeling experiment after pre-treating with a broad-spectrum inhibitor of periplasmic remodeling, imipenem. We observed similar patterns of fluorescence (Fig. S6) Taken together, our data demonstrate that peptidoglycan assembly continues in the absence of DivIVA but in a disorganized fashion.

**Figure 3.**
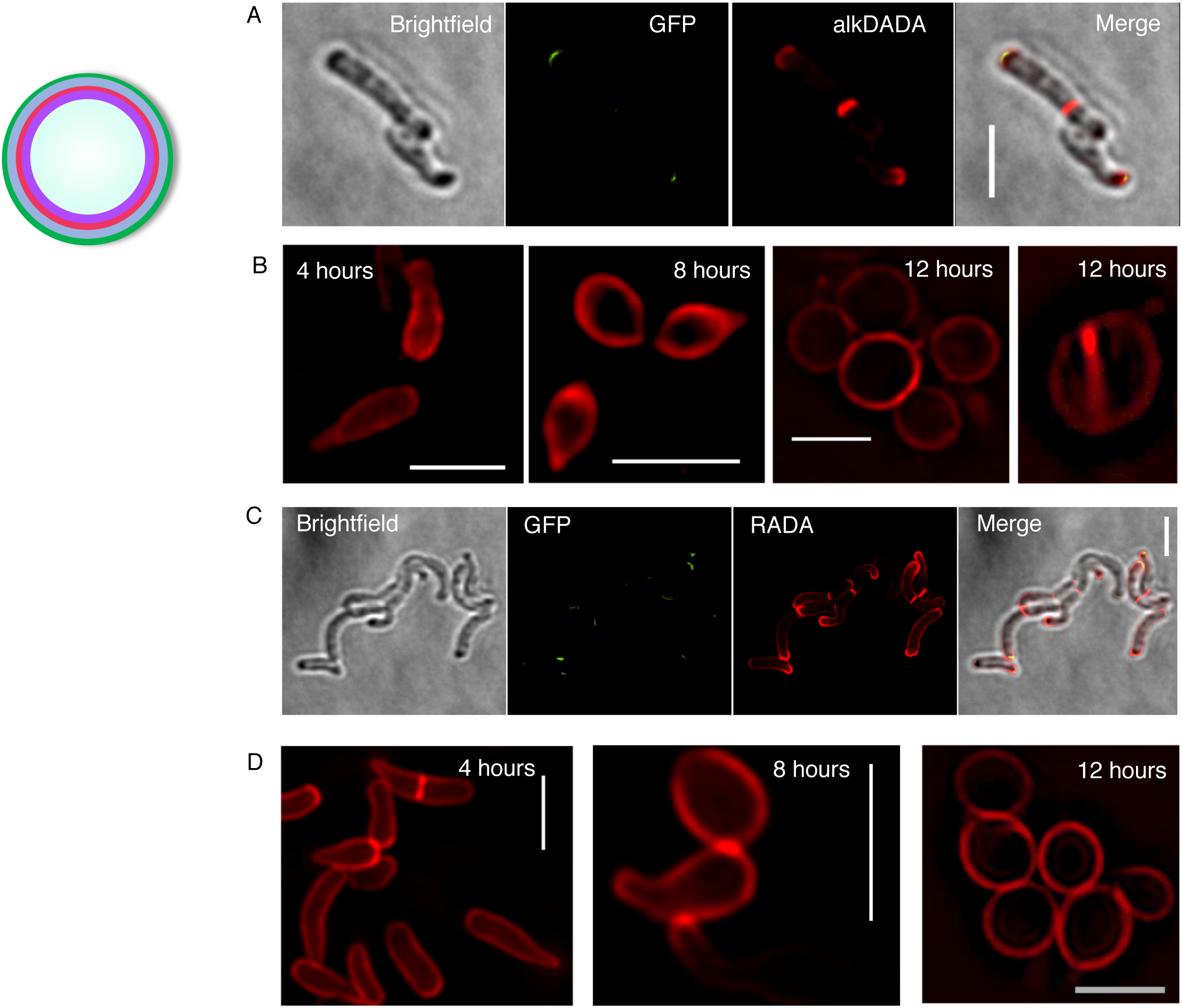
Peptidoglycan assembly is delocalized but persistent in the absence of DivIVA. DivIVA-eGFP *M. smegmatis* before (A) or during (B) ATC incubation was labeled with alkDADA for the final 15 minutes of ATC incubation time. Labeling was detected by CuAAC with picolyl azide-TAMRA. Scale bar, 5 µm. DivIVA-eGFP *M. smegmatis* before (C) or during (D) ATC incubation was labeled with RADA for the final 15 minutes of ATC incubation time. Scale bar, 5 µm. Exposure, (A) 1s (B) 2s (C) 500ms (D) 4 hrs: 400ms, 8 hrs: 600ms, 12 hrs: 400ms.

Assembly of the three, covalently-bound layers of the mycobacterial envelope—peptidoglycan, arabinogalactan and the mycomembrane—is likely to be spatially coincident. In support, the cytoplasmic enzymes that mediate arabinogalactan and mycomembrane synthesis are enriched at the mycobacterial cell poles [Meniche et al. 2014; Hayashi et al. 2016; Carel et al. 2014], as is metabolic labeling by trehalose monomycolate precursors that incorporate into the mycomembrane [Foley et al. 2016]. DivIVA physically interacts with enzymes required for early steps of mycolic acid precursor synthesis [Meniche et al. 2014; Xu et al. 2014]. To test whether DivIVA regulates the location of mycomembrane assembly, we labeled cells in the presence or absence of ATC with OalkTMM, a trehalose monomycolate probe that is primarily incorporated into arabinogalactan mycolates [Foley et al. 2016], and detected by CuAAC ligation to an azidefluorophore. In the absence of ATC, fluorescent signal derived from OalkTMM was again localized primarily to the septa and DivIVA-eGFP-marked poles, with limited sidewall labeling (Fig. 4A). In the presence of ATC, OalkTMM labeling was delocalized in a manner similar to alkDADA and RADA (Fig. 4B). As with peptidoglycan, assembly of the mycomembrane continues in the absence of DivIVA but is delocalized.

### Spatial and temporal coincidence of DivIVA localization, concentrated peptidoglycan assembly and rod extrusion from spherical M. smegmatis

Our data suggest that mycobacterial cell envelope assembly is persistent but disorganized in the absence of DivIVA (Fig. 3B, 3D, 4B). DivIVA-depleted cells are also unable to maintain their shape (Fig. 2D, 2E) [Kang et al. 2008; Nguyen et al. 2007; Meniche et al. 2014], suggesting that localized synthesis may support morphogenesis in these organisms. We wondered whether DivIVA might contribute to *de novo* morphogenesis by focusing envelope biogenesis. *E. coli* gradually resculpt rod shape following MreB inhibition or lysozyme or antibiotic treatment over rounds of division and dispersed elongation [Billings et al. 2014; Ranjit et al. 2017; Cambre et al. 2015]. In mycobacteria, however, rod-like protrusions appear to emerge directly from reverting spheroplasts or DivIVA-depleted spheres [Udou et al. 1982] (Fig. 2F). We hypothesized that the difference in outgrowth phenotype might be related to the ability of DivIVA to concentrate zones of peptidoglycan synthesis. We depleted DivIVA-eGFP for 16 hours until spheres were generated. We then washed away ATC and labeled the cells with alkDADA at the end of a 10 hour recovery period. The tips of the nascent protrusions were marked with alkDADA labeling, which in turn colocalized with DivIVA-eGFP foci (Fig. 5A). When outlines of recovering cells were used to generate relative fluorescence intensity profiles as they correspond to cell geometry, we found that parts of the cell that were still spherical had little or no visible eGFP and dimmer, more homogenous alkDADA-derived fluorescence (Fig. 5B). Additionally, we found that the intensity peaks for both channels were spatially correlative. Our data are consistent with a model in which DivIVA helps to establish and maintain shape in mycobacteria by nucleating cell envelope synthesis.

**Figure 4.**
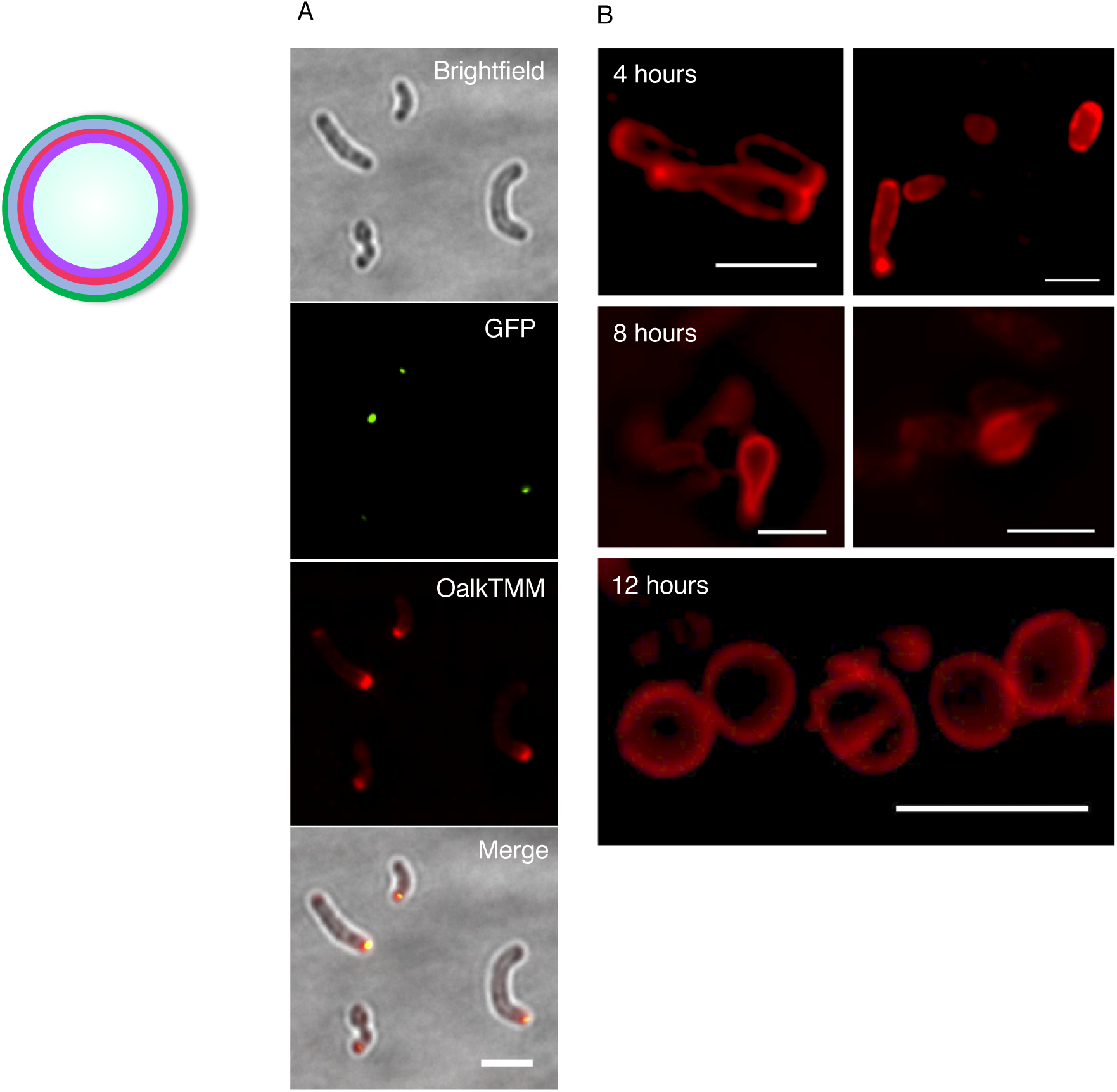
Mycomembrane assembly is delocalized but persistent in the absence of DivIVA. DivIVA-eGFP *M. smegmatis* before (A) or during (B) ATC incubation was labeled with OalkTMM for the final 15 minutes of ATC incubation time. Labeling was detected by CuAAC with azide-545. Scale bar, 5 µm. Exposure, 1s.

**Figure 5.**
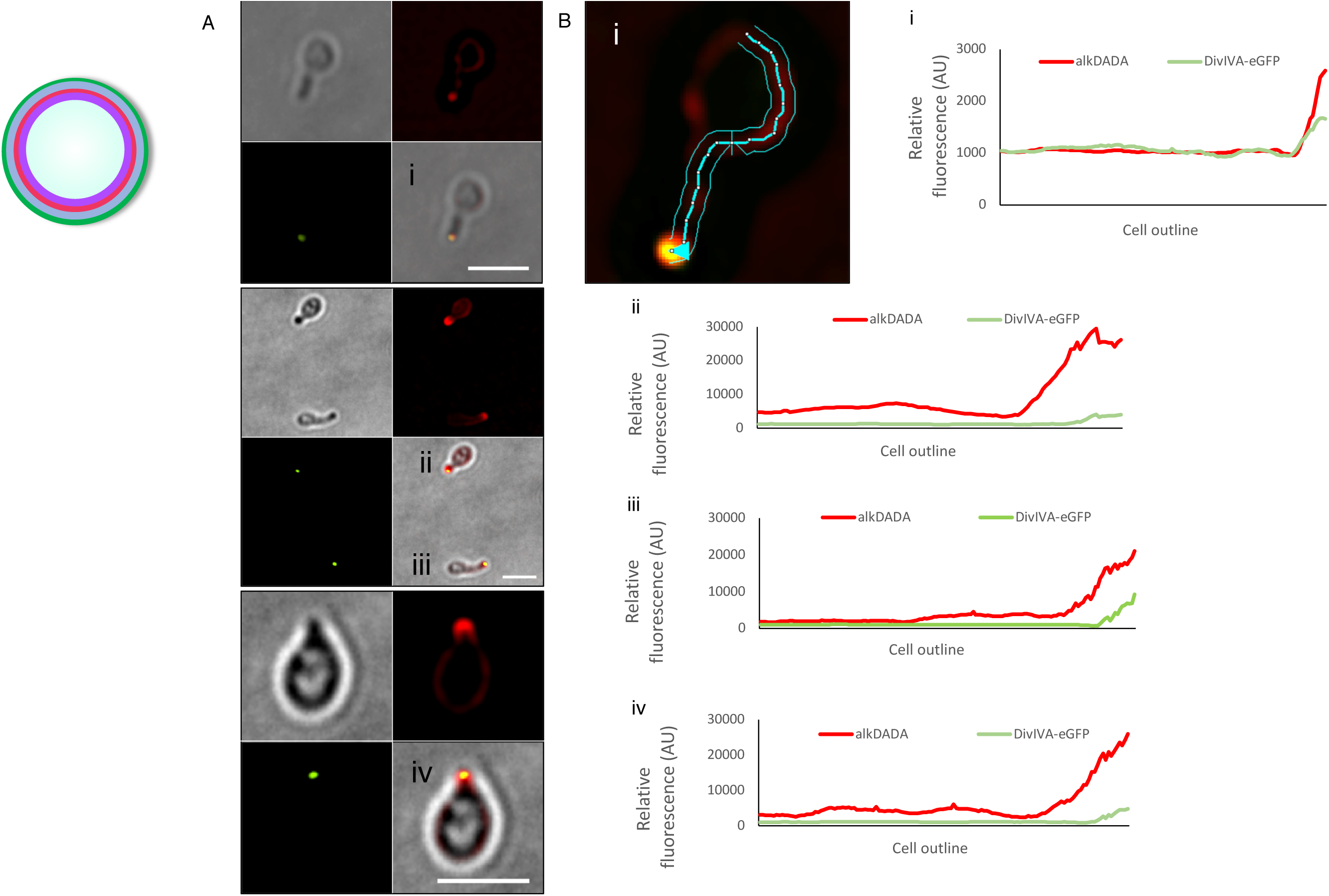
Correlation between rod outgrowth, site of DivIVA-eGFP focus and localized peptidoglycan assembly. (A) DivIVA-eGFP *M. smegmatis* were incubated in ATC for 16 hours and labeled 10 hours post-washout with alkDADA for the final 15 minutes of ATC incubation time. Labeling was detected by CuAAC with picolyl azide-TAMRA. Scale bar, 5 µm. (B) Lateral cell outlines were used to generate relative mean intensity profiles for eGFP and TAMRA signal as a function of individual cell geometry, using fluorescence of 258 nm on either side of the drawn outline. Intensity profile analysis applied to cells i, ii, iii, iv from (A). Results are representative of 12 cells.

## Summary of Results

The ability of a bacterium to readily regenerate its shape suggests that morphology is not simply a state to which the cell converges, but rather is hardwired [Billings et al. 2014]. Mycobacteria are evolutionarily distant and differ in both envelope composition and elongation mode. By visualizing the sphere-to-rod transition in *M. smegmatis*, we demonstrate here that programmed morphogenesis is a conserved trait. While MreB-containing species and *M. smegmatis* establish and propagate rod shape by cytoskeleton-controlled patterning of cell wall peptidoglycan, they do so via distinct mechanisms. MreB directs and is directed by peptidoglycan synthesis across a broad swath of the *E. coli* cell [Cabeen and Jacobs-Wagner 2010; Errington 2015; Carballido-Lopez and Errington 2003; Eun et al. 2015; Wagstaff and Lowe 2018]. By contrast, we show that DivIVA focuses cell envelope assembly in narrow, polar regions of both normal, rod-shaped mycobacteria and in rod-like protrusions from spherical cells. How does DivIVA control cell envelope synthesis? In the absence of the protein, we find that envelope assembly continues yet is disorganized. DivIVA repletion reverses this phenotype. Our data support a model in which both continuous and *de novo* polar envelope assembly depend on DivIVA-mediated concentration of the synthetic enzymes and/or their substrates and products [Meniche et al. 2014].

## Acknowledgments

We are grateful to Dr. Kadamba “Sundaram” Papavinasasundaram and Dr. Christopher Sassetti, University of Massachusetts Medical School, for helpful discussions and for providing the DivIVA-eGFP and RFP-DivIVA *M. smegmatis* strains. We also thank Dr. Benjamin Swarts, Central Michigan University, for OalkTMM and Dr. Eric Rubin, Harvard School of Public Health, for RADA. MSS is supported by NIH U01 CA221230 and NIH DP2 AI138238. ESM is supported by NIH T32 GM008515 administered to the Chemistry Biology Interface Program at the University of Massachusetts Amherst. SIM and time-lapse imaging were performed in the Light Microscopy Facility and Nikon Center of Excellence at the Institute for Applied Life Sciences, University of Massachusetts Amherst with support from the Massachusetts Life Science Center.

## Materials and Methods

### Generation of cell wall-deficient spheroplasts

Spheroplasts were generated by a modifying a protocol from [Xu et al. 2017; Udou et al. 1982; Udou et al. 1983]. Wildtype and DivIVA-eGFP *M. smegmatis* [Meniche et al. 2014] were grown to stationary phase in 7H9 medium (Difco) and back-diluted for overnight growth to reach OD_600_ = ~0.8. Glycine was added for a final concentration of 1.2% and cells were incubated shaking at 37°C for 24 hours. Cells were then washed once with a sucrose-MgCl_2_-maleic acid (SMM) buffer and centrifuged for 5 minutes at 4,000xg. Cells were resuspended in 7H9 that was prepared with SMM buffer in place of H_2_O. Glycine was added at a final concentration of 1.2% and lysozyme resuspended in SMM buffer was added at a final concentration of 50 µg/mL. Following incubation at 37^˚^C for 24 hours, spheroplasted cells were either imaged, labeled, or washed in SMM buffer and resuspended in fresh 7H9-SMM for recovery.

### Generation of DivIVA-depleted spheres

DivIVA-GFP *M. smegmatis* [Meniche et al. 2014] were grown to stationary phase and back-diluted for overnight growth to reach OD_600_ = ~0.8. Anhydrotetracycline (ATC; Sigma) was added to cultures at a final concentration of 50 ng/mL and cells were incubated shaking at 37°C for 4, 8, 12, 16 hours.

### Peptidoglycan and mycomembrane labeling

DivIVA-eGFP *M. smegmatis* [Meniche et al. 2014] were grown to stationary phase and back-diluted for overnight growth to reach OD_600_ = ~0.8. Cultures were incubated in the presence or absence of ATC for 4, 8 or 12 hours. Envelope precursor probes alkDADA (2 mM; custom synthesized by Albert Einstein College of Medicine Chemical Synthesis Core Facility), OalkTMM (50 µM; gift of Dr. Ben Swarts) or RADA (20 µM; gift of Dr. Eric Rubin) were added to the cultures for the final 15 minutes of incubation. Cells were washed once in cold PBS and fixed in 2% formaldehyde for 10 minutes. Cells were washed a second time in PBS then subjected to CuAAC with azide-545 or picolyl azide-TAMRA (Click Chemistry Tools) as described [Siegrist et al., 2013; Siegrist et al., 2015]. 12 hour DivIVA-depleted cells were also treated with 5 μg/mL clavulanate and 1 μg/mL imipenem for 30 min, then labeled with alkDADA for 15 min.

### Viability Staining

DivIVA-depleted cells were incubated with propidium iodide at a final concentration of 4 µM for 30 minutes at room temperature in the dark. As a control, non-depleted cells were treated with 70% isopropanol (168 of 169 stained nonviable), and live control was a population of non-depleted cells (7 of 67 stained nonviable).

### SIM Imaging and analysis

Images were acquired by Nikon Eclipse Ti N-SIM E microscope equipped with a Hamamatsu Orca Flash 4.0 camera with a numerical aperture of 1.49. Images were taken at 400 millisecond – 2 second exposure and were reconstructed and quantitated on NIS Elements. Cells were imaged on 1% agarose pads.

### Imaging and morphological analysis of spheroplasts

Recovering spheroplasts were placed on 1% agarose pads. Images were acquired on Nikon Eclipse E600. Manual quantification of cell morphology was scored blindly and processed using Fiji.

### Time-lapse imaging of recovery from DivIVA depletion

Following 16 hours of ATC incubation, DivIVA-eGFP *M. smegmatis* were washed in SMM buffer and resuspended in 7H9. 200 µL of concentrated cell suspension was plated onto 35 mm dishes [Joyce et al. 2011], liquid was aspirated 30 minutes later, and adherent cells were covered with a top layer of 0.7% agarose LB. Cells were incubated at 37°C for 8 hours and then placed in an ibidi heating system on a Nikon Eclipse Ti microscope with 100X DIC objective and Hamamatsu Orca Flash 4.0 camera, numerical aperture of 1.49. Time-lapse imaging was comprised of DIC image taken every 8 minutes, and fluorescence images taken every 40 minutes to reduce phototoxicity.

## Supplementary Material Legends

**Figure S1. Loss of peptidoglycan labeling in misshapen and spherical regions of cells during and after spheroplasting procedure**. *M. smegmatis* were labeled overnight with RADA and imaged prior to spheroplasting, (A), after 24 hours of glycine incubation, (B), and after completion of lysozyme digestion, (C). Scale bars, 5 µm.

**Figure S2. DivIVA foci persist in the absence of visible negative curvature.** Spheroplasts were generated from *M. smegmatis* expressing either DivIVA-eGFP, (A), or RFP-DivIVA, (B). Scale bars, 5 µm.

**Figure S3: Growth curves of DivIVA-eGFP M. smegmatis during ATC-induced depletion and following washout.** DivIVA-depleted cells grow more slowly during depletion and recovery.

Figure S4: Growth curves of wildtype *M. smegmatis* and DivVIA-eGFP *M. smegmatis*.

**Figure S5: Viability and rod protrusion upon DivIVA repletion.** (A) Ratio of spheres that exhibited protruding growth poles after ATC washout between 10-24 hours of recovery (n=52). (B) Ratio of DivIVA-depleted spheres that stained viable or nonviable when treated with propidium iodide (n=94).

**Figure S6: alkDADA labeling of DivIVA-depleted cells upon inhibition of periplasmic remodeling.** After 12 hours of DivIVA depletion, cells were treated with 5 μg/mL clavulanate and 1 μg/mL imipenem (2X minimal inhibitory concentration) for 30 min, then labeled with alkDADA for 15 min. Labeling was detected by CuAAC with picolyl azide-TAMRA. Scale bar, 5 µm.

**Movie S1: DivIVA-eGFP and rod protrusion from a sphere are spatiotemporally correlated.** Time-lapse microscopy tracking of a cell recovering from 16 hours DivIVA-eGFP depletion, starting 14 hours post-ATC removal, representative of 21 cells.

## References

Aldridge BB, Fernandez-Suarez M, Heller D, Ambravaneswaran V, Irimia D, Toner M and Fortune SM. 2012. Asymmetry and aging of mycobacterial cells lead to variable growth and antibiotic susceptibility. Science 335(6064):100-4.

Billings G, Ouzounov N, Ursell T, Desmarais SM, Shaevitz J, Gitai Z and Huang KC. 2014. De novo morphogenesis in L-forms via geometric control of cell growth. Mol Microbiol 93(5):883-96.

Birdsell DC and Cota-Robles EH. 1967. Production and ultrastructure of lysozyme and ethylenediaminetetraacetate-lysozyme spheroplasts of *Escherichia coli*. J Bacteriol 93(1):427-37.

Botella H, Yang G, Ouerfelli O, Ehrt S, Nathan CF and Vaubourgeix J. 2017. Distinct Spatiotemporal Dynamics of Peptidoglycan Synthesis between *Mycobacterium smegmatis* and Mycobacterium tuberculosis. MBio 8(5).

Cabeen MT and Jacobs-Wagner C. 2010. The bacterial cytoskeleton. Annu Rev Genet 44:365-92.

Cambre A, Zimmermann M, Sauer U, Vivijs B, Cenens W, Michiels CW, Aertsen A, Loessner MJ, Noben JP, Ayala JA et al. 2015. Metabolite profiling and peptidoglycan analysis of transient cell wall-deficient bacteria in a new *Escherichia coli* model system. Environ Microbiol 17(5):1586-99.

Carballido-Lopez R and Errington J. 2003. A dynamic bacterial cytoskeleton. Trends Cell Biol 13(11):577-83.

Carel C, Nukdee K, Cantaloube S, Bonne M, Diagne CT, Laval F, Daffe M and Zerbib D. 2014. Mycobacterium tuberculosis proteins involved in mycolic acid synthesis and transport localize dynamically to the old growing pole and septum. PLoS One 9(5):e97148.

Cha JH and Stewart GC. 1997. The divIVA minicell locus of Bacillus subtilis. J Bacteriol 179(5):1671-83.

Daniel RA and Errington J. 2003. Control of cell morphogenesis in bacteria: two distinct ways to make a rod-shaped cell. Cell 113(6):767-76.

de Pedro MA, Quintela JC, Holtje JV and Schwarz H. 1997. Murein segregation in *Escherichia coli*. J Bacteriol 179(9):2823-34.

Doi M, Wachi M, Ishino F, Tomioka S, Ito M, Sakagami Y, Suzuki A and Matsuhashi M. 1988. Determinations of the DNA sequence of the mreB gene and of the gene products of the mre region that function in formation of the rod shape of *Escherichia coli* cells. J Bacteriol 170(10):4619-24.

Dominguez-Escobar J, Chastanet A, Crevenna AH, Fromion V, Wedlich-Soldner R and Carballido-Lopez R. 2011. Processive movement of MreB-associated cell wall biosynthetic complexes in bacteria. Science 333(6039):225-8.

Drees B, Brown C, Barrell BG and Bretscher A. 1995. Tropomyosin is essential in yeast, yet the TPM1 and TPM2 products perform distinct functions. J Cell Biol 128(3):383-92.

Errington J. 2015. Bacterial morphogenesis and the enigmatic MreB helix. Nat Rev Microbiol 13(4):241-8.

Eun YJ, Kapoor M, Hussain S and Garner EC. 2015. Bacterial Filament Systems: Toward Understanding Their Emergent Behavior and Cellular Functions. J Biol Chem 290(28):17181-9.

Flardh K. 2003. Essential role of DivIVA in polar growth and morphogenesis in Streptomyces coelicolor A3(2). Mol Microbiol 49(6):1523-36.

Foley HN, Stewart JA, Kavunja HW, Rundell SR and Swarts BM. 2016. Bioorthogonal Chemical Reporters for Selective In Situ Probing of Mycomembrane Components in Mycobacteria. Angew Chem Int Ed Engl 55(6):2053-7.

Foss MH, Eun YJ and Weibel DB. 2011. Chemical-biological studies of subcellular organization in bacteria. Biochemistry 50(36):7719-34.

Fura JM, Kearns D and Pires MM. 2015. D-Amino Acid Probes for Penicillin Binding Protein-based Bacterial Surface Labeling. J Biol Chem 290(51):30540-50.

Garner EC, Bernard R, Wang W, Zhuang X, Rudner DZ and Mitchison T. 2011. Coupled, circumferential motions of the cell wall synthesis machinery and MreB filaments in *B. subtilis*. Science 333(6039):222-5.

Gitai Z, Dye N and Shapiro L. 2004. An actin-like gene can determine cell polarity in bacteria. Proc Natl Acad Sci U S A 101(23):8643-8.

Gunning P, Weinberger R and Jeffrey P. 1997. Actin and tropomyosin isoforms in morphogenesis. Anat Embryol (Berl) 195(4):311-5.

Hamasha K, Sahana MB, Jani C, Nyayapathy S, Kang CM and Rehse SJ. 2010. The effect of Wag31 phosphorylation on the cells and the cell envelope fraction of wild-type and conditional mutants of *Mycobacterium smegmatis* studied by visible-wavelength Raman spectroscopy. Biochem Biophys Res Commun 391(1):664-8.

Hayashi JM, Richardson K, Melzer ES, Sandler SJ, Aldridge BB, Siegrist MS and Morita YS. 2018. Stress-Induced Reorganization of the Mycobacterial Membrane Domain. MBio 9(1).

Hayashi JM, Luo CY, Mayfield JA, Hsu T, Fukuda T, Walfield AL, Giffen SR, Leszyk JD, Baer CE, Bennion OT et al. 2016. Spatially distinct and metabolically active membrane domain in mycobacteria. Proc Natl Acad Sci U S A 113(19):5400-5.

Hempel AM, Wang SB, Letek M, Gil JA and Flardh K. 2008. Assemblies of DivIVA mark sites for hyphal branching and can establish new zones of cell wall growth in *Streptomyces coelicolor*. J Bacteriol 190(22):7579-83.

Huang KC and Ramamurthi KS. 2010. Macromolecules that prefer their membranes curvy. Mol Microbiol 76(4):822-32.

Hussain S, Wivagg CN, Szwedziak P, Wong F, Schaefer K, Izore T, Renner LD, Holmes MJ, Sun Y, Bisson-Filho AW et al. 2018. MreB filaments align along greatest principal membrane curvature to orient cell wall synthesis. Elife 7.

Jani C, Eoh H, Lee JJ, Hamasha K, Sahana MB, Han JS, Nyayapathy S, Lee JY, Suh JW, Lee SH et al. 2010. Regulation of polar peptidoglycan biosynthesis by Wag31 phosphorylation in mycobacteria. BMC Microbiol 10:327.

Jankute M, Cox JA, Harrison J and Besra GS. 2015. Assembly of the Mycobacterial Cell Wall. Annu Rev Microbiol 69:405-23.

Jones LJ, Carballido-Lopez R and Errington J. 2001. Control of cell shape in bacteria: helical, actin-like filaments in *Bacillus subtilis*. Cell 104(6):913-22.

Joyce G, Robertson BD and Williams KJ. 2011. A modified agar pad method for mycobacterial live-cell imaging. BMC Res Notes 4:73.

Joyce G, Williams KJ, Robb M, Noens E, Tizzano B, Shahrezaei V and Robertson BD. 2012. Cell division site placement and asymmetric growth in mycobacteria. PLoS One 7(9):e44582.

Kang CM, Nyayapathy S, Lee JY, Suh JW and Husson RN. 2008. Wag31, a homologue of the cell division protein DivIVA, regulates growth, morphology and polar cell wall synthesis in mycobacteria. Microbiology 154(Pt 3):725-35.

Kawai Y, Mercier R and Errington J. 2014. Bacterial cell morphogenesis does not require a preexisting template structure. Curr Biol 24(8):863-7.

Kuru E, Tekkam S, Hall E, Brun YV and Van Nieuwenhze MS. 2015. Synthesis of fluorescent D-amino acids and their use for probing peptidoglycan synthesis and bacterial growth in situ. Nat Protoc 10(1):33-52.

Kuru E, Hughes HV, Brown PJ, Hall E, Tekkam S, Cava F, de Pedro MA, Brun YV and VanNieuwenhze MS. 2012. In Situ probing of newly synthesized peptidoglycan in live bacteria with fluorescent D-amino acids. Angew Chem Int Ed Engl 51(50):12519-23.

Lebar MD, May JM, Meeske AJ, Leiman SA, Lupoli TJ, Tsukamoto H, Losick R, Rudner DZ, Walker S and Kahne D. 2014. Reconstitution of peptidoglycan cross-linking leads to improved fluorescent probes of cell wall synthesis. J Am Chem Soc 136(31):10874-7.

Lenarcic R, Halbedel S, Visser L, Shaw M, Wu LJ, Errington J, Marenduzzo D and Hamoen LW. 2009. Localisation of DivIVA by targeting to negatively curved membranes. EMBO J 28(15):2272-82.

Liang H, DeMeester KE, Hou CW, Parent MA, Caplan JL and Grimes CL. 2017. Metabolic labelling of the carbohydrate core in bacterial peptidoglycan and its applications. Nat Commun 8:15015.

Liechti GW, Kuru E, Hall E, Kalinda A, Brun YV, VanNieuwenhze M and Maurelli AT. 2014. A new metabolic cell-wall labelling method reveals peptidoglycan in Chlamydia trachomatis. Nature 506(7489):507-10.

Margolin W. 2009. Sculpting the bacterial cell. Curr Biol 19(17):R812-22.

Marston AL and Errington J. 1999. Selection of the midcell division site in *Bacillus subtilis* through MinD-dependent polar localization and activation of MinC. Mol Microbiol 33(1):84-96.

Meniche X, Otten R, Siegrist MS, Baer CE, Murphy KC, Bertozzi CR and Sassetti CM. 2014. Subpolar addition of new cell wall is directed by DivIVA in mycobacteria. Proc Natl Acad Sci U S A 111(31):E3243-51.

Nguyen L, Scherr N, Gatfield J, Walburger A, Pieters J and Thompson CJ. 2007. Antigen 84, an effector of pleiomorphism in *Mycobacterium smegmatis*. J Bacteriol 189(21):7896-910.

Onoda T, Oshima A, Nakano S and Matsuno A. 1987. Morphology, growth and reversion in a stable L-form of *Escherichia coli* K12. J Gen Microbiol 133(3):527-34.

Plocinska R, Purushotham G, Sarva K, Vadrevu IS, Pandeeti EV, Arora N, Plocinski P, Madiraju MV and Rajagopalan M. 2012. Septal localization of the *Mycobacterium tuberculosis* MtrB sensor kinase promotes MtrA regulon expression. J Biol Chem 287(28):23887-99.

Plocinski P, Martinez L, Sarva K, Plocinska R, Madiraju M and Rajagopalan M. 2013. Mycobacterium tuberculosis CwsA overproduction modulates cell division and cell wall synthesis. Tuberculosis (Edinb) 93 Suppl:S21-7.

Plocinski P, Ziolkiewicz M, Kiran M, Vadrevu SI, Nguyen HB, Hugonnet J, Veckerle C, Arthur M, Dziadek J, Cross TA et al. 2011. Characterization of CrgA, a new partner of the *Mycobacterium tuberculosis* peptidoglycan polymerization complexes. J Bacteriol 193(13):3246-56.

Ramamurthi KS and Losick R. 2009. Negative membrane curvature as a cue for subcellular localization of a bacterial protein. Proc Natl Acad Sci U S A 106(32):13541-5.

Ramirez-Arcos S, Liao M, Marthaler S, Rigden M and Dillon JA. 2005. *Enterococcus faecalis* divIVA: an essential gene involved in cell division, cell growth and chromosome segregation. Microbiology 151(Pt 5):1381-93.

Ramos A, Honrubia MP, Valbuena N, Vaquera J, Mateos LM and Gil JA. 2003. Involvement of DivIVA in the morphology of the rod-shaped actinomycete *Brevibacterium lactofermentum*. Microbiology 149(Pt 12):3531-42.

Ranjit DK and Young KD. 2016. Colanic Acid Intermediates Prevent De Novo Shape Recovery of *Escherichia coli* Spheroplasts, Calling into Question Biological Roles Previously Attributed to Colanic Acid. J Bacteriol 198(8):1230-40.

Ranjit DK, Jorgenson MA and Young KD. 2017. PBP1B Glycosyltransferase and Transpeptidase Activities Play Different Essential Roles during the De Novo Regeneration of Rod Morphology in Escherichia coli. J Bacteriol 199(7).

Rastogi N and Venkitasubramanian TA. 1979. Preparation of protoplasts and whole cell ghosts from *Mycobacterium smegmatis*. J Gen Microbiol 115(2):517-21.

Rego EH, Audette RE and Rubin EJ. 2017. Deletion of a mycobacterial divisome factor collapses single-cell phenotypic heterogeneity. Nature 546(7656):153-57.

Santi I, Dhar N, Bousbaine D, Wakamoto Y and McKinney JD. 2013. Single-cell dynamics of the chromosome replication and cell division cycles in mycobacteria. Nat Commun 4:2470.

Sarkar S, Libby EA, Pidgeon SE, Dworkin J and Pires MM. 2016. In Vivo Probe of Lipid II-Interacting Proteins. Angew Chem Int Ed Engl 55(29):8401-4.

Scheffers DJ and Pinho MG. 2005. Bacterial cell wall synthesis: new insights from localization studies. Microbiol Mol Biol Rev 69(4):585-607.

Siegrist MS, Swarts BM, Fox DM, Lim SA and Bertozzi CR. 2015. Illumination of growth, division and secretion by metabolic labeling of the bacterial cell surface. FEMS Microbiol Rev 39(2):184-202.

Siegrist MS, Whiteside S, Jewett JC, Aditham A, Cava F and Bertozzi CR. 2013. (D)-Amino acid chemical reporters reveal peptidoglycan dynamics of an intracellular pathogen. ACS Chem Biol 8(3):500-5.

Singh B, Nitharwal RG, Ramesh M, Pettersson BM, Kirsebom LA and Dasgupta S. 2013. Asymmetric growth and division in Mycobacterium spp.: compensatory mechanisms for non-medial septa. Mol Microbiol 88(1):64-76.

Singh V, Dhar N, Pato J, Kolly GS, Kordulakova J, Forbak M, Evans JC, Szekely R, Rybniker J, Palcekova Z et al. 2017. Identification of aminopyrimidine-sulfonamides as potent modulators of Wag31-mediated cell elongation in mycobacteria. Mol Microbiol 103(1):13-25.

Swarts BM, Holsclaw CM, Jewett JC, Alber M, Fox DM, Siegrist MS, Leary JA, Kalscheuer R and Bertozzi CR. 2012. Probing the mycobacterial trehalome with bioorthogonal chemistry. J Am Chem Soc 134(39):16123-6.

Takacs CN, Poggio S, Charbon G, Pucheault M, Vollmer W and Jacobs-Wagner C. 2010. MreB drives de novo rod morphogenesis in *Caulobacter crescentus* via remodeling of the cell wall. J Bacteriol 192(6):1671-84.

Thanky NR, Young DB and Robertson BD. 2007. Unusual features of the cell cycle in mycobacteria: polar-restricted growth and the snapping-model of cell division. Tuberculosis (Edinb) 87(3):231-6.

Thomaides HB, Freeman M, El Karoui M and Errington J. 2001. Division site selection protein DivIVA of *Bacillus subtilis* has a second distinct function in chromosome segregation during sporulation. Genes Dev 15(13):1662-73.

Tiyanont K, Doan T, Lazarus MB, Fang X, Rudner DZ and Walker S. 2006. Imaging peptidoglycan biosynthesis in *Bacillus subtilis* with fluorescent antibiotics. Proc Natl Acad Sci U S A 103(29):11033-8.

Udou T, Ogawa M and Mizuguchi Y. 1982. Spheroplast formation of *Mycobacterium smegmatis* and morphological aspects of their reversion to the bacillary form. J Bacteriol 151(2):1035-9.

Udou T, Ogawa M and Mizuguchi Y. 1983. An improved method for the preparation of mycobacterial spheroplasts and the mechanism involved in the reversion to bacillary form: electron microscopic and physiological study. Can J Microbiol 29(1):60-8.

van den Ent F, Amos LA and Lowe J. 2001. Prokaryotic origin of the actin cytoskeleton. Nature 413(6851):39-44.

van Teeffelen S, Wang S, Furchtgott L, Huang KC, Wingreen NS, Shaevitz JW and Gitai Z. 2011. The bacterial actin MreB rotates, and rotation depends on cell-wall assembly. Proc Natl Acad Sci U S A 108(38):15822-7.

Wachi M, Doi M, Okada Y and Matsuhashi M. 1989. New mre genes mreC and mreD, responsible for formation of the rod shape of *Escherichia coli* cells. J Bacteriol 171(12):6511-6.

Wagstaff J and Lowe J. 2018. Prokaryotic cytoskeletons: protein filaments organizing small cells. Nat Rev Microbiol doi:10.1038/nrmicro.2017.153.

Wang S, Furchtgott L, Huang KC and Shaevitz JW. 2012. Helical insertion of peptidoglycan produces chiral ordering of the bacterial cell wall. Proc Natl Acad Sci U S A 109(10):E595-604.

Wei JR, Krishnamoorthy V, Murphy K, Kim JH, Schnappinger D, Alber T, Sassetti CM, Rhee KY and Rubin EJ. 2011. Depletion of antibiotic targets has widely varying effects on growth. Proc Natl Acad Sci U S A 108(10):4176-81.

Xu WX, Zhang L, Mai JT, Peng RC, Yang EZ, Peng C and Wang HH. 2014. The Wag31 protein interacts with AccA3 and coordinates cell wall lipid permeability and lipophilic drug resistance in *Mycobacterium smegmatis*. Biochem Biophys Res Commun 448(3):255-60.

Xu Z, Meshcheryakov VA, Poce G and Chng SS. 2017. MmpL3 is the flippase for mycolic acids in mycobacteria. Proc Natl Acad Sci U S A 114(30):7993-98.

Yabu K and Takahashi S. 1977. Protoplast formation of selected *Mycobacterium smegmatis* mutants by lysozyme in combination with methionine. J Bacteriol 129(3):1628-31.

Young KD. 2006. The selective value of bacterial shape. Microbiol Mol Biol Rev 70(3):660-703.

